# Antimicrobial Resistance Profile and *mcr-1* Gene Detection in *Salmonella* Isolates from Poultry in Bangladesh: Molecular and Bioinformatics Characterization

**DOI:** 10.1101/2020.03.27.012948

**Authors:** Md Bashir Uddin, S M Bayejed Hossain, Mahmudul Hasan, Mohammad Nurul Alam, Mita Debnath, Ruhena Begum, Sawrab Roy, Ahmed Harun-Al-Rashid, Md. Shahidur Rahman Chowdhury, Md. Mahfujur Rahman, Md. Mukter Hossain, Mohammed Yousuf Elahi Chowdhury, Syed Sayeem Uddin Ahmed

## Abstract

Antimicrobial resistance gene *mcr-1* has been disseminated globally since its first discovery in Southern China in late 2015. However, the *mcr*-1 gene had not been identified previously in *Salmonella* isolates from poultry in Bangladesh. Here, we aimed to explore antimicrobial resistance gene *mcr-1* in *Salmonella* isolates. Eighty two *Salmonella* isolates were isolated and characterized from suspected poultry specimens received from different zones of the country. A phenotypic disc diffusion assay with 15 antimicrobial agents was performed following CLSI standard. The disk diffusion assay showed that, all of the isolates presented high resistance to colistin (92.68%), oxytetracycline (86.59%), co-trimoxazole (76.83%), ciprofloxacin (73.17%) and enrofloxacin (65.85%). Further, randomly selected 10 *Salmonella* isolates were analyzed by polymerase chain reaction (PCR) targeting genus-specific *invA* and antimicrobial (colistin) resistance *mcr*-1 genes. Five were confirmed for the presence of the *mcr-1* gene belonging to *Salmonella* spp. Further, sequencing followed by phylogenetic analysis revealed divergent evolutionary relation between the LptA and MCR proteins rendering them resistant to colistin. Three-dimensional homology structures of MCR-1 proteins were constructed and verified using different bioinformatics tools. Moreover, molecular docking interactions suggested that, MCR-1 and LptA share a similar substrate binding cavity which could be validated for the functional analysis. The results represent here is the first molecular and *in silico* analysis of colistin resistance *mcr-1* gene of *Salmonella* in poultry in Bangladesh, which may emphasize the importance of the study on antibiotic resistance genes requiring for national monitoring and strategic surveillance in the country.

## Introduction

Poultry (mainly broiler and layer) farming is an important avenue in fostering agricultural growth and has becoming a major contributing sector for potential income generation and poverty alleviation in Bangladesh (1, 2). Eggs and poultry meats are most acceptable and largely consumed animal products to meet dietary nutritional requirements throughout the country (3, 4). In spite of significant improvement, there is a potential threat of diseases due to bacterial infections that can result a huge economic loss in this sub-sector (5, 6). Among them, infections with *Salmonella* spp. are the most commonly reported problem in poultry that cause food borne illness to human and remain as persistent threat to both human and animal health (6–8). Globally, 94 million human affected cases were estimated due to *Salmonella* spp. leading to 155,000 deaths every year (9, 10).

Salmonellosis is endemic in nature causing morbidity and mortality in poultry. It is very significant by virtue of the fact that, it can be transmitted vertically from parent to offspring; this makes its control a challenge. Although vaccination and good hygiene practices are most effective ways to prevent salmonellosis (11), antibiotics are extensively using either as a growth promoter or prophylaxis and therapeutics in poultry industry of Bangladesh (12, 13). Indeed, the widespread misapplication and nonjudicious use of antimicrobial drugs in poultry settings, culminating the development of antimicrobial resistant pathogens like *Salmonella* (Gyles, 2008, Cantas et al., 2013; Antunes et al., 2016).

Antimicrobial resistant *Salmonella* of poultry can harbor as a major risk and vehicle for dissemination of these pathogens to humans (16, 17). Standard culture, biochemical and serological methods are usually employed for isolation and identification of *Salmonella* species. However, the *invA* encoding invasion gene, commonly involved in bacterial virulence, is accountable and routinely used for the detection of *Salmonella* spp. (18). Moreover, the *invA* sequences are distinctive to the genus *Salmonella* and diagnosed by polymerase chain reaction (PCR) is a preferred diagnostic method due to its reliable sensitivity, specificity and detection speed (19, 20).

Of note, poultry is usually incriminated in outbreaks of human Salmonellosis. Therefore, the detection of *Salmonella* species in poultry production chain particularly at the farm level is an issue of great concern. Furthermore, the resistance of some *Salmonella* serotype to multiple antibiotics (15) makes the study on antibiotic susceptibility profile and its antimicrobial resistance gene, a great priority (21). So far, *mcr-1* positive *Enterobacteriaceae* (MCRPE) has been found in animal, food, human and environment in over 25 countries across 4 continents (22–25). As far as literature mining is concerned from PubMed search regarding poultry in Bangladesh, no data was found on the molecular characterization and antimicrobial resistance gene detection in *Salmonella*.. Therefore, the present study has undertaken to detect colistin resistance *mcr-1* gene for the first time in *Salmonella* isolates and its associated drug resistance pattern of commonly used antibiotics.

In this study, morphological and biochemical techniques were generated and phenotypic and genotypic characteristics of isolates were explored using antimicrobial susceptibility testing, PCR, nucleotide analysis, bioinformatics and structural modeling of bacterial genetics. The phylogenetic relationships between local isolates and published data sets from different corner of the world were analyzed. Additionally, molecular docking of phosphatidylethanolamine substrate with MCR-1 and LptA were investigated. Focus has been given on antimicrobial resistance *mcr-1* gene involved in multidrug leading to colistin resistance.

## Materials and methods

### Ethical standards

The research has been conducted in accordance with the Institutional Ethics Committee of Kazi Farms Group, Dhaka and Sylhet Agricultural University (SAU), Sylhet, Bangladesh.

### Study area and sampling

The study was conducted from January to June 2019 at popular poultry zones of Bangladesh: Gazipur, Narsingdi, Tangail and Brahmanbaria (**Figure 1**). Samples of dead and sick birds were collected and transferred to the customer service lab at Kazi Farms Group, Gazipur, Bangladesh. Postmortem was conducted immediate after receiving samples with their anamnesis and clinical information in accordance with the standard guidelines by a veterinarian. During postmortem, liver and intestinal samples were collected and immediately sent for further analysis. For sick live birds, blood was collected for serum separation before the postmortem examination.

**Figure 1:**
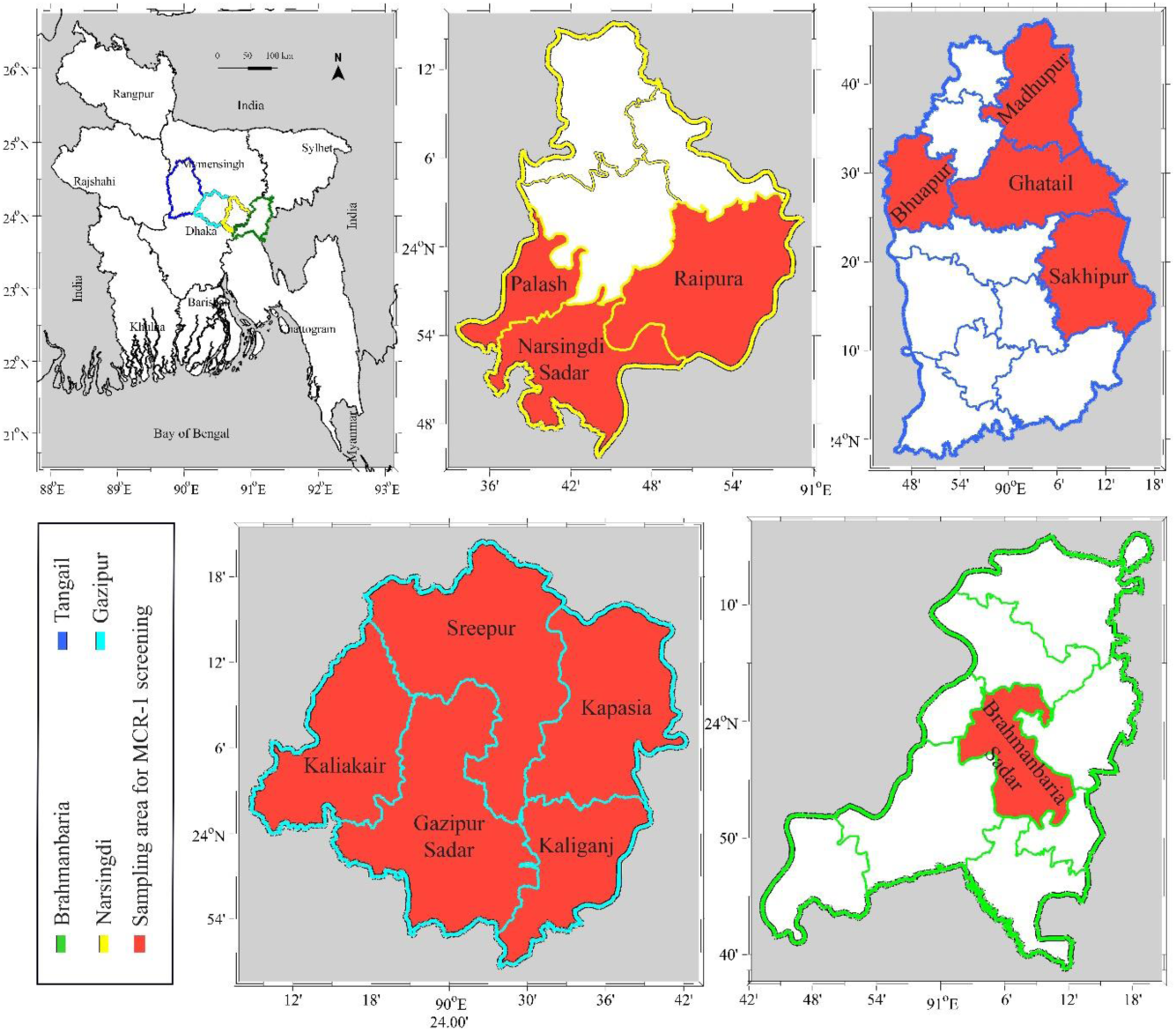
A map of Bangladesh showing locations of the sampling sites under selected districts. The areas where the *mcr-1* gene has been screened are highlighted in red.

### Isolation and biochemical identification of bacterial isolates

Before isolation of *Salmonella* spp, samples were initially screened out by Rapid Serum Plate Agglutination Test (RSPAT; ID Vet, France) followed by clinical and postmortem findings. Liver and intestinal samples from 100 suspected samples were subjected to a pre-enrichment step by combining with 225 mL Buffered Peptone Water (BPW) in a ratio of 10 fold dilutions and incubated at 37°C for 24 hours (h). For *Salmonella* specific pre-enrichment, culture were further transferred to Modified Semi-solid Rappaport Vassiliadis (MSRV; Hi Media, India) and Tetrathionate Broth (TTB; Hi Media, India) consecutively and incubated at 42°C for 24 h. Following enrichment, a loop of enriched broth was initially streaked on Xylose-Lysine-Desoxycholate (XLD; Hi Media, India) agar and colonies (single pinkish) were streaked on *Salmonella-Shigella* (SS) agar, incubated at 37°C for 24 h. Salmonella colonies were identified by physical and biochemical properties using Gram’s stain, catalase and indole tests as previously described (Dashti et al., 2009; Sobur et al., 2019). For further confirmation, 4-6 suspected *Salmonella* colonies from each samples were tested biochemically by dilution streaking and stab onto Triple Sugar Iron (TSI) agar (Merck, Germany) and incubation at 37°C for 16-24 h (28, 29).

### Antimicrobial susceptibility testing

*Salmonella* isolates were examined for phenotypic antibiogram using 15 antimicrobial agents by Kirby–Bauer disk diffusion method as previously described (30). In a brief, isolates were grown on Mueller-Hinton (MH) agar (Hi Media, India) and incubated for 16-18 h at 37°C. McFarland 0.5 standards were maintained for culture suspension of individual isolate. Discs were placed on the agar surface using a sterile forceps and incubated at 37°C for 18 hours. The tested results were interpreted by measuring the zones of inhibition and scored as sensitive, intermediate and resistant according to the CLSI (CLSI, 2019) (**Table1**). Isolates that were found resistant against at least 3 classes of antibiotics considered as multidrug resistance (MDR) *Salmonella* spp (**Table 2**) (32).

**Table 1:**
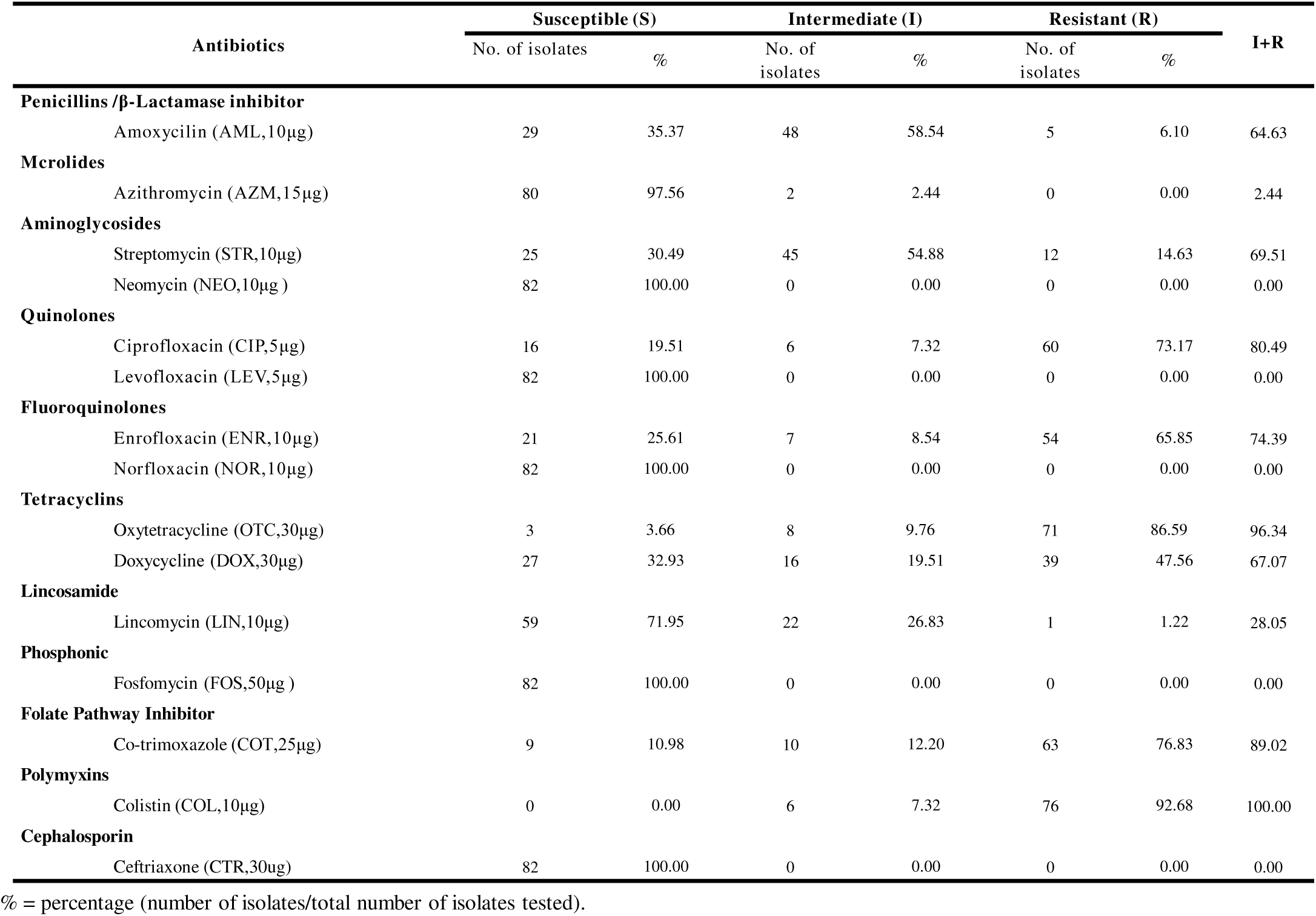
Antimicrobial susceptibility pattern of *Salmonella* isolates (n=82).

**Table 2:** Multidrug resistant patterns among the *mcr-1* positive *Salmonella* spp isolated from poultry farms in Bangladesh

| *Salmonella*<br>isolates | Multidrug resistant patterns | No. of antibiotic classes to<br>which the tested isolates<br>exhibits resistance ($n = 7$) |
| --- | --- | --- |
| **S-6** | COL+OTC+COT+CIP+STR | 5 |
| **S-7** | COL+OTC+COT+CIP | 4 |
| **S-8** | COL+OTC+AZM+CIP+STR | 5 |
| **S-9** | COL+OTC+COT+CIP | 4 |
| **S-10** | COL+OTC+COT | 3 |

COL = Colistin, OXT = Oxytetracycline, AZM= Azithromycin, COT= Co-trimoxazole, CTR=Ceftriaxone, CIP = Ciprofloxacin, STR = Streptomycin.
Resistant if, Colistin: $\leq 10$ ; Tetracycline: $\leq 11$ ; Azithromycin: $\leq 12$ ; Ceftriaxone: $\leq 14$ ; Ciprofloxacin: $\leq 20$ ; Streptomycin: $\leq 11$ .
Multidrug resistant patterns was determined according to the recommendations by CLSI zone diameter interpretation criteria (nearest whole mm).

### Extraction of Bacterial genomic DNA

Among 82 *Salmonella* isolates, 10 isolates were randomly subjected to molecular characterization for the identification of resistance genes. For this, bacterial DNA was extracted by boiling-centrifugation method as described earlier (27, 33). In a brief, a loop full of overnight cultured bacterial suspension was transferred into 1.5 mL microcentrifuge tube and centrifuged at 13,000 rpm for 1 min. The supernatant was discarded and 1 mL of sterile ultrapure water was added and vortexed. The suspension was heated at 100°C for 8∼10 min in a heating block and then immediately cooled on ice for 5 min. Cell debris from the cell lysates were pelleted by centrifugation at 13,000 rpm for 1 min and remaining supernatant was used as DNA templates for PCR assays.

### Polymerase Chain Reaction (PCR) and gel electrophoresis

Extracted DNA was subjected to PCR for the initial confirmation of *Salmonella* isolates using specific primers (**Table 3**) targeting *invA* gene with the expected amplicon size of 100 bp. A PCR assay was further performed with confirmed 10 *Salmonella* isolates to detect the presence of antimicrobial resistance gene *mcr-1*. In this case, 2 targeting primer sets (**Table 4**) were designed for covering frame reading of antimicrobial (colistin) resistance gene *mcr-1* with the allocated 2 amplicon sizes (1197 bp and 799 bp). Both PCR assays were performed in 20 mL reaction mixture containing 5 uM of each forward and reverse primer and 5 μL of extracted genomic DNA as template. *Salmonella* positive plasmid (5 μL) used as positive controls (PC) and sterile molecular grade water was used as negative controls (NC) to detect cross-contamination during DNA purification and PCR. To optimize PCR reaction, lambda DNA amplification (1000 bp) was used as internal control (IC). PCR was performed using PCR thermocycler (Bio-Rad, United States); conditions for amplification of *invA* and *mcr-*1genes were listed in **Supplementary Table 1 & 2**. The amplified products were visualized by gel electrophoresis using 1.5% agarose gel and viewed under UV transilluminator in Gel Documentation System (Bio-Rad, United States). For *mcr-1* gene, PCR products of 2 primer sets were purified (Addbio Inc., product code: 10078, South Korea) and used as direct sequencing. Two sequences of each sample was assembled for covering frame reading of *mcr-1* gene and checked with BLAST and annotated to GenBank. In the case of *invA* gene, 10 representative amplicons of *invA* gene fragments were sequenced to confirm the identity of *invA* gene using BLAST (**Supplementary File 1**). DNA sequencing was done by commercial sequencing company SolGent (Daejeon, Republic of Korea).

**Table 3:**
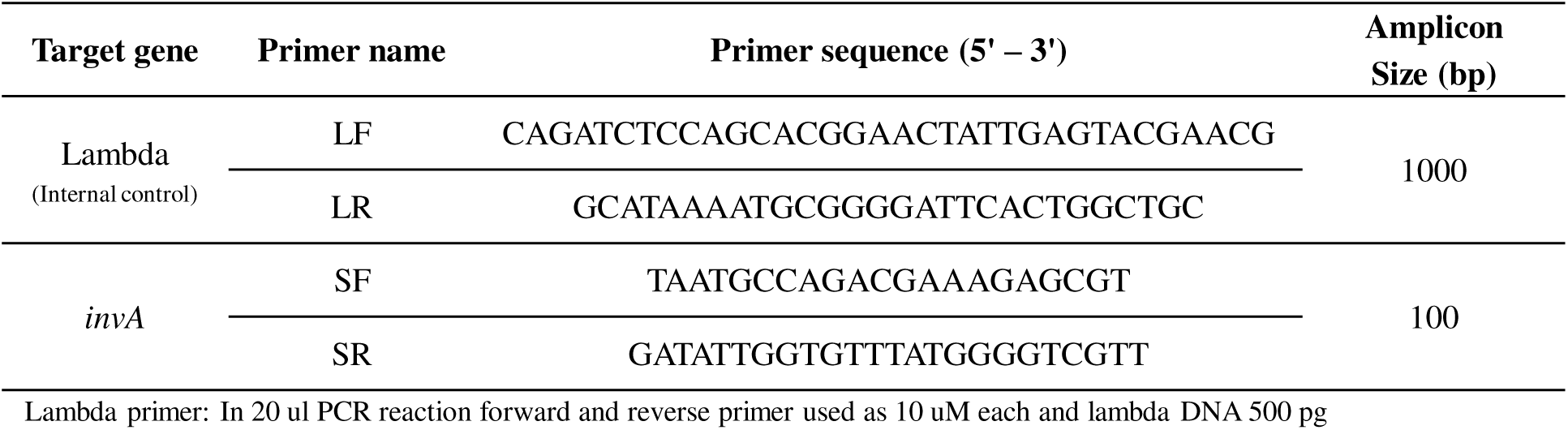
Primer sequence for *invA* gene detection in *Salmonella* isolates

**Table 4:**
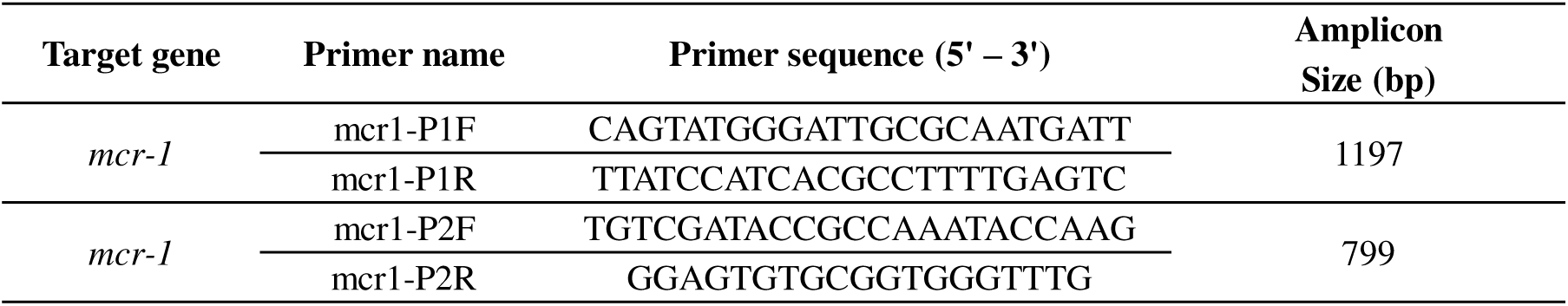
Primer sequence for *mcr-1* gene detection in *Salmonella* isolates

### Sequence acquisition, multiple sequence alignment and phylogenetic analysis

*BLASTp* search (https://blast.ncbi.nlm.nih.gov/Blast.cgi) was employed to retrieve the homologous sequences of the MCR-1 and MCR-1 like proteins from the NCBI database using six SAUVM MCR-1 proteins translated from the *mcr-1* genes of *Salmonella spp.* Sequences were carefully categorized into MCR-1 and MCR-1 Like proteins of *Salmonella, E. coIi* strains, strains containing LptA (formerly named EptA) and others (**Supplementary File 2 and 3**). Multiple sequence alignment of SAUVM*mcr-1* proteins and retrieved *mcr-1* of *Salmonella* species were performed using T-Coffee with default parameters (34). Maximum Likelihood Method of MEGA X (35) was employed to construct a phylogenetic tree using aligned sequences of MCR-1 from ClustalW (36). Results were validated using 500 bootstrap replicates.

### Transmembrane topology analysis, structural modelling, refinement and validation

To predict the transmembrane helices of MCR-1 proteins, TMHMM server (http://www.cbs.dtu.dk/services/TMHMM/) was used with standard parameters. The topology was given as the position of the transmembrane helices differentiated by ‘i’ and ‘o’ when the loop is on the inside and outside respectively (37). Three dimensional (3D) modelling of SAUVM MCR-1 proteins were performed by I-TASSER which functions by identifying structure templates from the Protein Data Bank (PDB) library. I-TASSER simulations generate large ensemble of structural conformations based on the pair-wise structure similarity. The confidence of each model is quantitatively measured by C-score (38). To enhance the accuracy of the predicted structures, refinement was performed using ModRefiner (39) followed by FG-MD refinement server (40, 41). Finally, the refined structures were also validated using Verified 3D (42, 43), ERRAT (44) and Ramachandran Plot Assessment server (RAMPAGE) (45, 46).

### Molecular docking of PE substrate with *mcr-1* and LptA

The chemical structure of Phosphatidylethanolamine (PE) (ZINC identification number [ID]: ZINC32837869) was sampled from the ZINC database (47) while the 3D structure of LptA (PDB ID: 5FGN; Organism: Neisseria meningitidis), the best template of SAUVM MCR-1 were retrieved from the RCSB Protein Data Bank (PDB) server (48). Binding interactions of the PE in the MCR-1 LptA was investigated by molecular docking using Autodock Vina algorithm in PyRx software (49). OpenBabel (version 2.3.1) was used to convert the output PDBQT files in PDB format. PyMol and Discovery Studio software were used to optimize and visualize the protein structures and ligand binding interaction patterns (50, 51).

### Nucleotide Sequence Accession Number

The sequences of five *mcr-1* genes of *Salmonella* spp. isolated from poultry were deposited into the GenBank database with the Accession No. MN873694, MN873695, MN873696, MN873697 and MN873698 for SAUVM_S6, SAUVM_S7, SAUVM_S8, SAUVM_S9 and SAUVM_10, respectively.

## Results

### Confirmation of *Salmonella* spp. and their antimicrobial susceptibility

*Salmonella* isolates were confirmed by morphological (Gram’s stain), cultural (MacConkey, XLD and S-S agar media) and biochemical (catalase, indole and TSI agar slants test) characteristics as previously described (29). A total of 82 *Salmonella* isolates were differentiated and confirmed following their morphological and biochemical properties. Out of 82 isolates, 10 were further confirmed for the presence of *invA* virulence genes, which is accountable for salmonellosis. PCR (using specific primer listed in **Table 3)** was conducted as a confirmatory detection tool and found 10 out of 10 (100%) suspected *Salmonella* isolates were *invA* gene (100 bp) positive. Amplified DNA fragment of 100 base pairs were considered as positive for *Salmonella* isolates (**Figure 2**). All the *Salmonella* isolates (n=82) were subjected to antibiotic resistance profiling to 15 antimicrobials (**Table 1**). In general, a considerable percentage of resistance was observed across the entire isolates in disk diffusion assay according to the standard of CLSI (31). Specifically, high resistances were found against colistin (92.68%), oxytetracycline (86.59%), co-trimoxazole (76.83%), ciprofloxacin (73.17%) and enrofloxacin (65.85%).Isolates were shown 100% susceptible to the Ceftriaxone (CTR, 30ug), Fosfomycin (FOS,50μg), Norfloxacin (NOR,10μg), Levofloxacin (LEV,5μg), Azithromycin (AZM,15μg), and Neomycin (NEO,10μg) antimicrobials. Only 1 isolate (1/82, 1.22%) was resistant to Lincomycin (LIN, 10μg). Other isolate intermediately resistant to Amoxycilin (AML, 10μg), Streptomycin (STR, 10μg) and Lincomycin (LIN, 10μg). Although, there were some variations, some antibiotic profiles were common among isolates. The multidrug resistant (MDR) patterns were also evaluated among *mcr-1* positive *Salmonella* spp against different antimicrobial classes. All the isolates (100%) showed MDR against 3 antimicrobial classes (colistin, oxytetracycline and ciprofloxacin) (**Table 2**).

**Figure 2:**
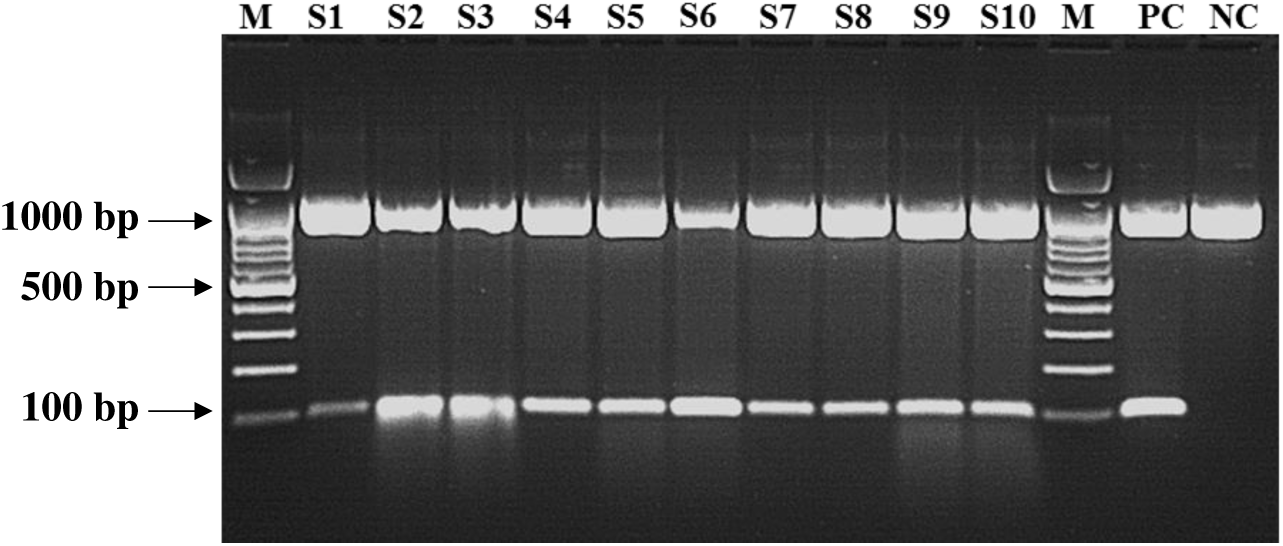
PCR amplification of *invA* gene of *Salmonella* isolates. In all isolates (S1∼S10), a fragment of 100 bp (*invA* gene) was detected. Lane M: DNA ladder. Lane PC: positive control for *invA* gene; Lane NC: negative control for *invA* gene; Lanes S1-S10, amplified gene of *invA* in the tested isolates. [S1∼10=*Salmonella* isolate 1∼10].

### Detection of colistin resistance *mcr-1* gene

*Salmonella* isolates were further evaluated for the presence of the *mcr-1* gene through PCR using *mcr-1* specific primers (**Table 4**). Five samples were likely to carry colistin resistance *mcr-1* among 10 randomly selected *Salmonella* isolates obtained from chickens (**Figure 3**). In Sanger sequencing demonstration, sequence found 100% identical with *mcr-1* gene accessed in the NCBI database and *mcr-1* gene described by Liu et al. (2016).

**Figure 3:**
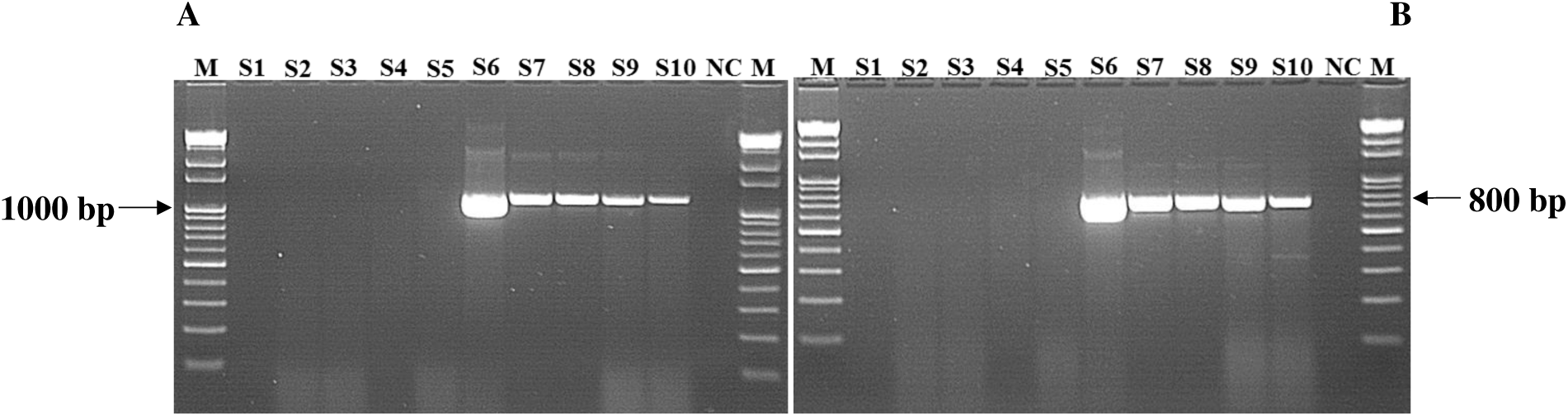
PCR amplification of antimicrobial (colistin) resistance *mcr-1* gene of *Salmonella* isolates. In *Salmonella* (S6∼S10) isolates a fragment of (A) 1197 bp and (B) 799 bp was detected. Lane M: DNA ladder. Lane NC: negative control for *mcr-1* gene. [S6∼10=*Salmonella* isolate 6∼10].

### Sequence acquisition, multiple sequence alignment and phylogenetic analysis

In order to analyze sequence similarities, phylogeny and structural insights of *mcr-1* gene products, the respective translated MCR-1 proteins were employed in different bioinformatics studies. A total of 52 homologous sequences of the MCR-1 and MCR-1 like proteins were retrieved from the NCBI database, while 44 sequences were employed to phylogenetic analysis including SAUVM-MCR-1 proteins. Again, 8 MCR-1 proteins of *Salmonella* spp were aligned with SAUVM-MCR-1 proteins for further analysis. The evolutionary relation inferred via phylogeny analysis has been given in **Figure 4**. The phylogenetic analysis revealed that, all of the MCR-1 and MCR-1 like proteins were distinctly categorized into 2 major groups; chromosomally-encoded LptA and plasmid encoded MCR types, indicating a divergent evolutionary relation between the LptA and MCR proteins.

**Figure 4:**
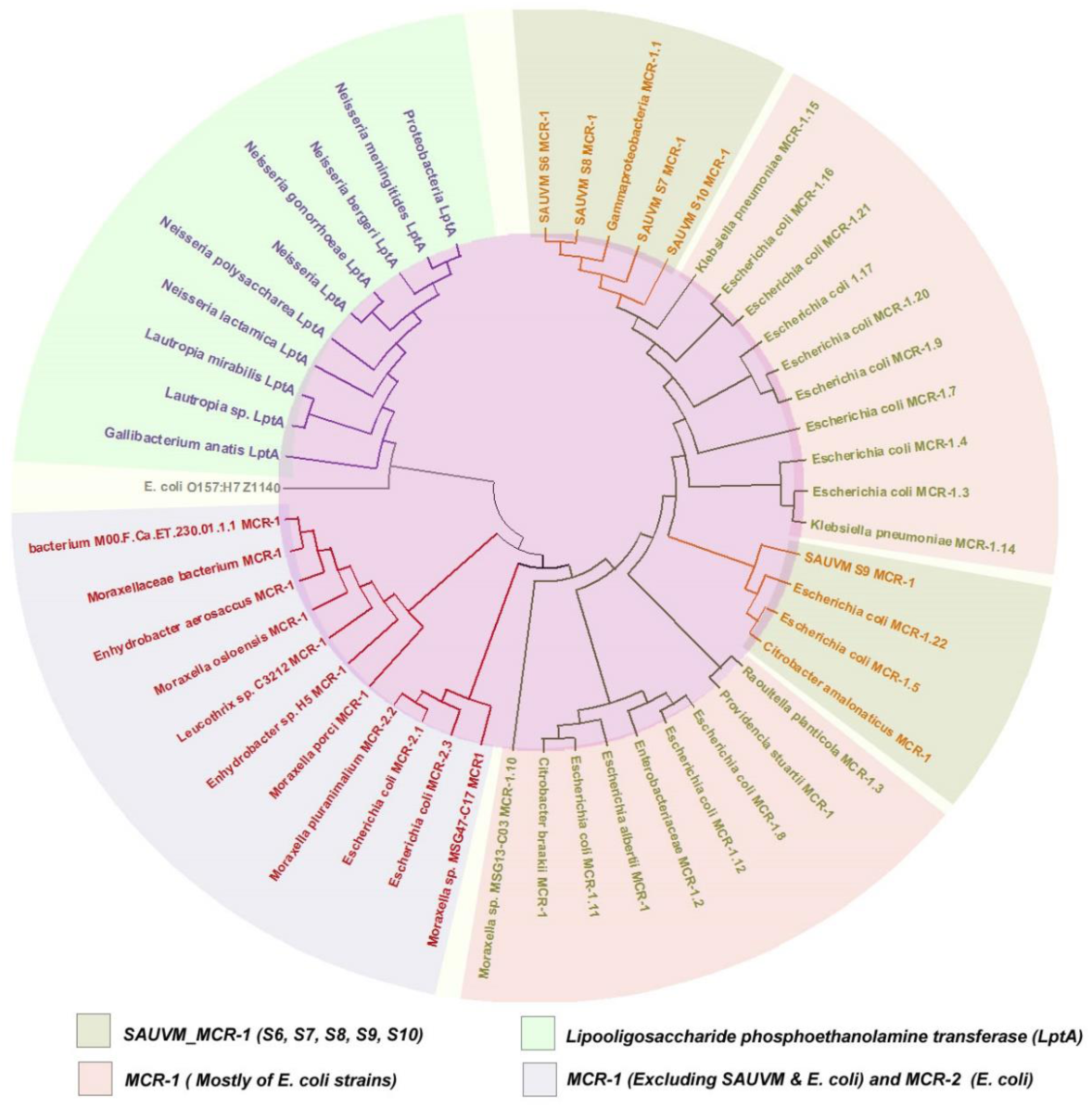
Phylogeny analysis showing ancestral origin and diversification of MCR-1 and MCR-1 like proteins. *BLASTp* search (https://blast.ncbi.nlm.nih.gov/Blast.cgi) was employed to retrieve the homologous sequences of the MCR-1 and MCR-1 like proteins from the NCBI database using amino acid sequences of six SAUVM-MCR-1 proteins. Sequences were carefully categorized into the MCR-1 and MCR-1 Like proteins of *Salmonella, E. coli* strains, strains containing LptA (formerly named EptA) and others. Maximum Likelihood Method of MEGA X was employed to construct a phylogenetic tree using aligned sequences of MCR-1 from CLUSTALW.

### Transmembrane topology analysis, structural modelling, refinement and validation

Prediction of transmembrane helices is of utmost importance in functional analysis of protein. Therefore, TMHMM server was applied for predicting transmembrane helices in mcr-1 genes of *Salmonella* isolates. TMHMM predicted that, there were five transmemebrane domain in the SAUVM-MCR-1 proteins; TMhelix1 (13-35), TMhelix 2 (50-72), TMhelix 3 (79-101), TMhelix 4 (123-145) and TMhelix 5 (158-180) which were spanned in the inner membrane region (**Figure 5**). The structure of SAUVM-MCR-1 proteins were modeled using I-TASSER server, where *N. meningitis* EptA (PDB ID 5FGN) acted as the structural template. SAUVM-MCR-1 proteins showed 35.4% (35.6%) identity to EptA, and their modelled structure possesses a coverage score of 96% compared with that of EptA. Refinement was performed to enrich the quality of predicted structures beyond the accuracy. After refinement Ramachandran plot analysis revealed that, 83.3% residues were in the favored, 12.4% residues in the allowed while only 4.3% residues were in the outlier region (**Figure 6**). Moreover, ERRAT showed 94.4% quality factor (**Supplementary Figure 1A and 1B**) and Verify3D suggested that, 94.74% of the residues had averaged 3D-1D score >= 0.2 (**Supplementary Figure 2**).

**Figure 5:**
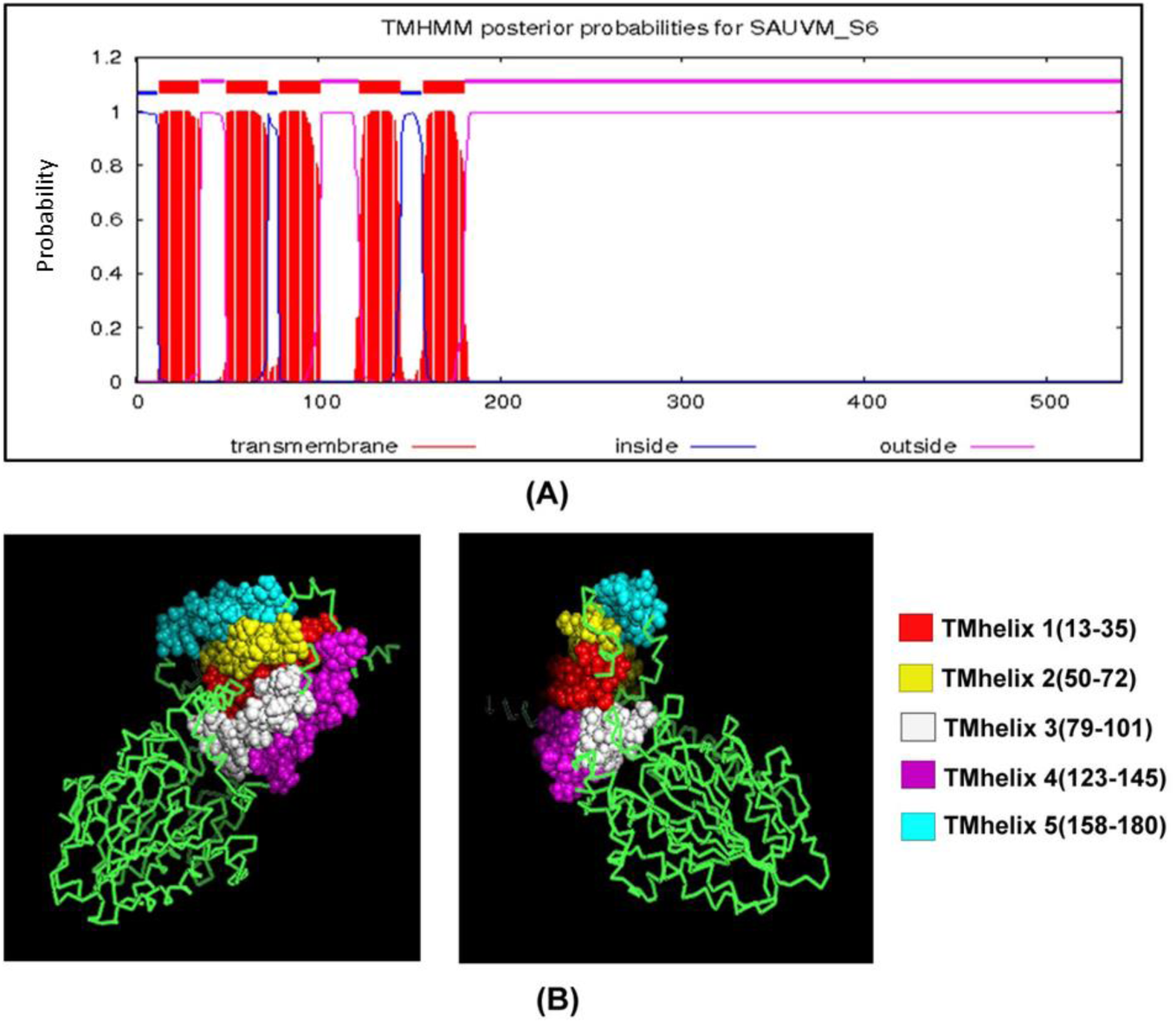
Transmembrane topology prediction of SAUVM-S6-MCR-1 protein. TMHMM server (http://www.cbs.dtu.dk/services/TMHMM/) was used to predict the (**A**) transmembrane helices of MCR-1 proteins. (**B**) The topology was given as the position of the transmembrane helices differentiated by ‘i’ and ‘o’ when the loop is on the inside and outside, respectively.

**Figure 6:**
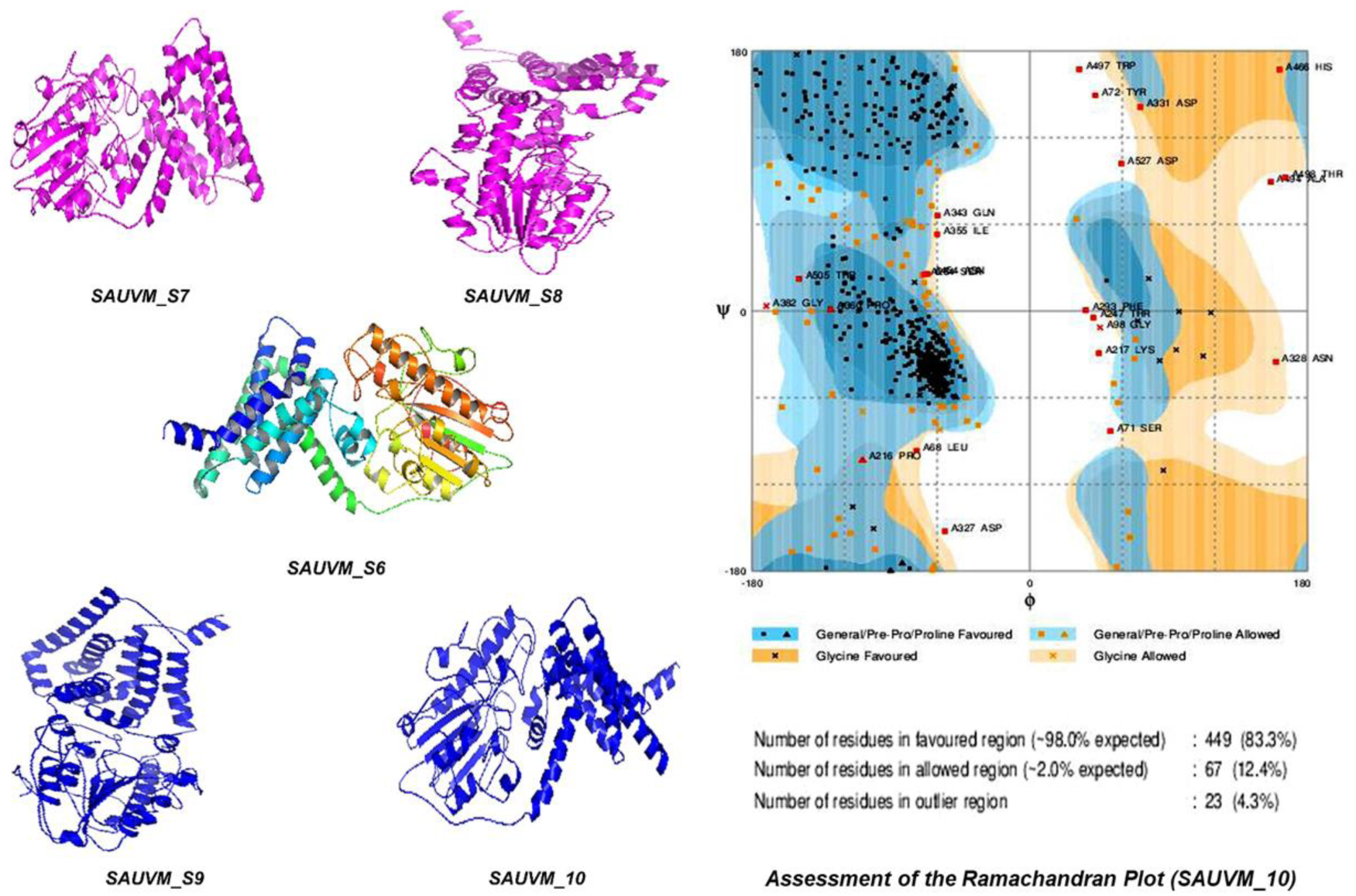
Modelled structures of SAUVM-MCR-1 proteins and Validation. (**A**) Three dimensional (3D) modelling of SAUVM-MCR-1 proteins (SAUVM_S6∼S10) were performed by I-TASSER which functions by identifying structure templates from the PDB library. The confidence of each model is quantitatively measured by C-score. From these models of MCR-1 proteins, SAUVM_S10 model was randomly selected, (**B**) analysed and structures validated with Ramachandran Plot Assessment server (RAMPAGE).

### Molecular docking of PE substrate with MCR-1 and LptA

The grid box was set to 82.0138A° x 82.7041A° x 82.471A° (x, y and z) with 1 A° spacing between the grid points, while other parameters were default. Though, molecular docking of PE substrate with SAUVM-MCR-1 and LptA generated five docking binding conformation for each but the binding pattern with lower energy had been selected (PE & MCR-1: −3.4kcal/mol and PE & LptA: −3.6 kcal/mol). Again, it was demonstrated that Leu 64, Tyr 179 and Phe 183 were the key interactive molecules in PE binding cavity of SAUVM MCR-1 whereas Ser 61, Tyr 174, Phe 181, Val 192 and Ser 194 were for LptA (**Figure 7**).

**Figure 7:**
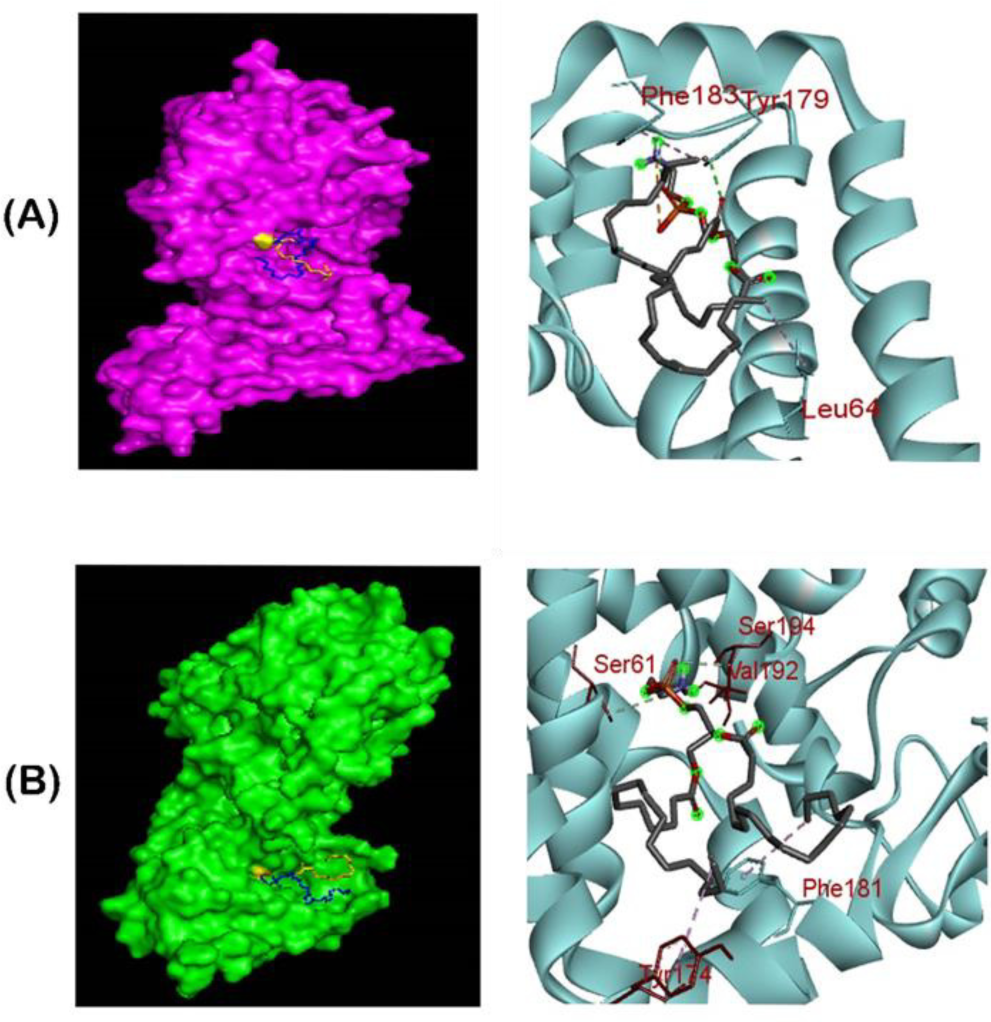
Ligand-binding interaction pattern of PE substrate with colistin resistance MCR-1 and LptA. The modeled ribbon structure for PE substrate with MCR-1 protein. The ribbon structure was given via PyMol software. In both (**A**) and (**B**) cases, the substrates tend to bind in the groove of MCR-1 and LptA mostly spanning from 175-195 region, in which Phe, Tyr and Ser residues were abundantly found in the substrate binding region for PE interaction. (**B**) Ligand-binding interaction revealed that both MCR-1 and LptA proteins exhibited similar localization of PE binding sites (SAUVM-MCR-1: Leu 64, Tyr 179, Phe 183; LptA: Ser 61, Tyr 174, Phe 181, val 192, Ser 194).

## Discussion

*Salmonella* is the primary and leading cause of food borne diseases globally; accounting for 78 million affected and 59 thousand deaths annually (52, 53). This is an endemic food borne disease in South Asian countries like Bangladesh (54, 55). A broad range of food stuff has been associated with such illness. Among them, food from animal sources, especially poultry is in main list. Due to its potential to cause enteric disease, the detection of *Salmonella* isolates in poultry is of great concern which are globally recognized as food borne zoonoses (52, 55, 56). The severity and length of such diseases could reduce by antimicrobial therapeutics in human (53) and poultry as well (57, 58). However, the rising of antimicrobial-resistant that is, antimicrobials commonly prescribed to treat *Salmonella* are losing its ability to stop growing or killing of *Salmonella* has become a significant public health issue now days (Islam and Shiraj-um-mahmuda, 2016; Ahmed et al., 2019 Wasyl et al., 2015; Iwamoto et al., 2017). As a result, standard treatment become ineffective, infections persist, and may increases chance of spreading to others (53, 61). Therefore, antimicrobial resistance is of great concern and challenging for any country. So far, Bangladesh requires baseline data on resistant bacteria like *Salmonella* and their molecular detection from various sources (e.g. poultry) to develop effective strategies against antimicrobial resistance and its hazards.

In the present study, to explore morphological, biochemical and molecular detection of antimicrobial resistant *Salmonella* and their genes in poultry, samples were obtained from both broiler and layer birds that were dead or clinically sick. Initial screening for infection has been done on the basis of clinical history, signs and postmortem findings. Among 100 suspected samples from different poultry zones of the country, 82 found positive for *Salmonella* spp. which was not unexpected as previous studies found that, even apparently healthy commercial poultry and their surrounding environment can carry *Salmonella* spp. in Bangladesh (62, 63).

Molecular methods were optimized for rapid identification of *Salmonella* isolates and its resistance genes in poultry using specific primers followed by nucleotide sequencing and phylogenetic analysis. The virulence of *Salmonella* is linked to a combination of various factors, for example, *invA* virulence gene in the inner membrane of bacteria that are necessary for invasion of epithelial cells (18, 19). In this study, detection of virulence *invA* gene (10 out of 10 isolates) in the isolated *Salmonella* indicates the pathogenic nature of these isolates.

Resistance of *Salmonella* spp. to antimicrobials is an emerging threat in developing as well as developed countries (7). It is therefore necessary to determine the resistance patterns of isolates to minimize resistance hazards. In this study, antibiotic susceptibility results showed 100% of the *Salmonella* isolates were resistant to colistin and oxytetracycline. This high resistance rates reflect widespread use of these antibiotics in animal feed and are consistent with other reports (25, 64). And rate of resistance to ciprofloxacin (80.49%) and enrofloxacinin (74.39%) deserves attention because *Salmonella* spp. resistance to these antibiotics may cause human infection too (58). However, previous studies reported that, *Salmonella* isolates were sensitive to ciprofloxacin (7, 62, 65). Similar results were observed in amoxicillin and doxycycline antimicrobials. These antimicrobials resistance against *Salmonella* isolates might designate the over use or abuse of these antibiotics (14, 15, 58). On the other hand, *Salmonella* isolates found to be susceptible to ceftriaxone, fosfomycin, neomycin, levofloxacin, norfloxacin and azithromycin which is in line with previous reports (64, 66). It may be because; these antibiotics are not commonly used for therapeutic purposes in veterinary medicine.. The present study represented that, 100% of the *mcr-1* positive isolates were MDR. Similar findings were reported on MDR in *Salmonella* isolates from Bangladesh and different parts of the world (29, 67–72). Due to the indiscriminate victimization of antimicrobial agents, MDR strains may apparently be occurred with high incidence in this area which is a serious concern for veterinary medicine and also for human health since direct transmission of resistant isolates from animals to humans has been confirmed (73).

Since early 1980s, colistin has been widely used in agricultural sector in China (25, 74), caused the initial emergence and spread of *mcr-1* worldwide (25, 68, 75, 76). In this study, an unexpected presence of (5 out 10 samples) colistin resistance *mcr-1* gene was detected in *Salmonella* spp. from poultry specimens (e.g. liver, intestine). This higher rate of *mcr-1* in chicken *Salmonella* isolates was surprising and suggested that *mcr-1* might already be widespread in food animals in Bangladesh. However, we do not have actual data on antimicrobial usage on the farms where the samples originated. Therefore, the presence of *mcr-1* gene may suggest frequent use of colistin and possibly other antimicrobials in the poultry industry in this region. A recent study has been reported 28% of poultry samples harbored *mcr-1* in China which has linked between human and animals (25). Subsequent study found an unexpected high prevalence (24.8%) of *mcr-1* in retail chicken meat samples in Netherlands (77). Following that work, investigations by other scientists have confirmed the presence of *mcr-1* in *Salmonella* isolates recovered from mussels and poultry (77–80). The high prevalence of the *mcr-1* gene in *Enterobacteriace* isolates of poultry is certainly concerning for a country like Bangladesh where antimicrobial use in both human and animals may be poorly regulated. Therefore, the recent emergence of colistin resistance has triggered an international review and recommendations for restrictions of colistin use in farm animals (68, 81).

While, detailed study on molecular mechanisms of antimicrobial resistance is lacking, we aimed to address *mcr-1* using integrative approaches ranging from nucleotide analysis, bioinformatics and structural modeling of bacterial genetics. The detection of new *mcr-1*-harbouring *Salmonella* isolates adds new knowledge to the newly-emerging issue of colistin resistance *mcr-1* genes. It furthering our understanding on homology, structure and validation of the *mcr*-*1* genes present in *Salmonella* isolates. It was reported that, colistin resistance *mcr*-*1* gene is present in a multidrug-resistant plasmid (82) and *mcr-1* genes of this study found somewhat similar to a recently-isolated plasmid from China (25). These facts imply that, multidrug resistant bacteria with colistin resistance will eventually evolve a fact that deserves close attention.

Thus, we are awfully interested in determining the multiple sequence alignments for *mcr-1* genes in *Salmonella* spp. isolated from poultry. In order to avoid hits from very closely related species, retrieved sequences of *Salmonella* species were excluded from the phylogeny study and those were only aligned with SAUVM *mcr-1* proteins. The multiple sequence alignments of SAUVM *mcr-1* proteins clearly indicated that, they belongs to the Mobilized Colistin Resistance (MCR) protein family with putative conserved sites [**Supplementary File 2**].

We experimentally validated the expression of colistin resistance *mcr-1* of *Salmonella* isolates (**Figure 3A and 3B**), suggesting the possibility of evolutionary path for the *mcr-1* genes. To address this concern, we conducted phylogenetic analyses. The phylogeny indicates a divergent evolutionary pattern between the LptA and MCR-1 including MCR-1 like proteins. The constructed phylogenetic tree provides information about ancestral origin and diversification of the MCR-1 proteins in different organism divided into chromosomally-encoded LptA and plasmid encoded MCR types indicating a divergent evolutionary relation between the LptA and MCR proteins (Figure 4). Further, MCR proteins group were divided into 2 apparent subgroups, one of which features MCR-1 proteins mostly of *E. coli* strains including SAUVM MCR-1 and the other one compromising small subclade of MCR-2 with MCR-1 proteins from diverse organisms. All of the SAUVM MCR-1 proteins were closely related to the *E. coli* MCR-1 sharing the position in the same clade. However, SAUVM MCR-1 and LptA fall into 2 separate subclades within the tree which indicated the low sequence identity, also reported by previous studies (83, 84). Despite the fact that, *E. coli* MCR-2 proteins were mostly aligned with MCR-1 proteins of non *E. coli* groups. Again, the Z1140 locus of *E. coli* O157:H7, a member of the PEA lipid a transferases lacking a role in colistin resistance apparently formed individual clade in the phylogeny which strengthened the findings. For understanding the structural insight, 3D homology modeling of five of SAUVM MCR-1 proteins were constructed using Neisseria meningitides LptA as structural template and five distinct transmembrane helices spanned in the inner membrane region were identified which was also reported by different studies (68, 84). Again, molecular interactions between Phosphatidylethanolamine (PE) substrate with MCR-1 and LptA had been investigated as colistin resistance proteins. MCR-1, MCR-2 and LptA were found to share similar PE lipid substrate-recognizing cavity. Ligand-binding interaction pattern of PE substrate with SAUVM-MCR-1 and LptA revealed that, both proteins exhibited similar localization of PE binding sites spanning from 175-195 region in which Phe, Tyr and Ser residues were abundantly found.

The data we present represents a first comprehensive glimpse of antimicrobial (colistin) resistance *mcr-1* genes among poultry originated *Salmonella* isolates in Bangladesh. This study provides further substantial evidence for the need of implementation of risk-management strategies and the need to review the extensive use of colistin in food animals for urgently advocated and implemented in this country.

## Acknowledgements

We thank customer service lab at Kazi Farms Group, Gazipur, Bangladesh for their technical support.

## Supplementary Files

**Supplementary File 1:** Amplicons of 100bp *invA* gene sequence of *Salmonella* isolates (S6∼S10).

**Supplementary File 2:** Retrieved sequences of MCR-1 and MCR-1 like proteins.

**Supplementary File 3:** The multiple sequence alignments of SAUVM-MCR-1 proteins with putative conserved sites of other *Salmonella* MCR-1.

**Supplementary File 4:** Supplementary Table S1, S2 and S3.

**Supplementary File 5:** Structure validation by ERRAT and Verify3D server.

